# NanoLoop: A deep learning framework leveraging Nanopore sequencing for chromatin loop prediction

**DOI:** 10.1101/2025.05.03.651998

**Authors:** Wenjie Huang, Li Tang, Matthew C. Hill, Yonghao Zhang, Jun Xie, Min Li

## Abstract

Chromatin loops play a crucial role in gene regulation and cellular function, providing key insights into understanding the three-dimensional structure of the genome and its impact on cellular homeostasis. Nanopore sequencing technology, with its advantages in simultaneously detecting sequences and methylation patterns, brings new opportunities for studying three-dimensional genome structures. We introduce NanoLoop, the first algorithmic framework attempting to predict genome-wide chromatin interactions using Nanopore data. In experiments across four human lymphoblastoid cell lines, NanoLoop demonstrated excellent predictive performance and cross-cell line generalization capabilities. We also discovered four distinct methylation patterns at loop anchors that influence histone modification levels and determine various loop types. NanoLoop further predicted previously uncharacterized long-range chromatin loops, highlighting DNA methylation’s role in three-dimensional genome regulation and providing new insights into the complex regulatory relationships between epigenetic modifications and three-dimensional genome organization.

## Introduction

DNA is not only the blueprint of life but also the foundation of the three-dimensional genomic architecture that governs cellular function and gene regulation ^1–3^. Chromatin loops are fundamental to organizing the genome’s spatial structure, enabling regulatory elements such as enhancers, promoters, and insulators to come into proximity with target genes. These loops facilitate precise control of transcriptional activity, influencing gene expression and maintaining cellular homeostasis. Understanding chromatin loops provides critical insights into the mechanisms of genome organization and its implications for development, disease, and cellular function ^4–7^.

With the continuous advancement of chromatin conformation capture (3C) technologies, methods such as Hi-C ^8^, ChIA-PET ^9^, and HiChIP/PLAC-Seq ^10,11^ have emerged, each offering distinct advantages in capturing chromatin interactions. These technologies have significantly propelled the development of three-dimensional genomics ^12–14^.

With the rapid advancement of artificial intelligence and deep learning technologies, algorithms for predicting chromatin loops based on Hi-C data have emerged ^15–18^. However, these algorithms often overlook the influence of epigenetic modifications on three-dimensional chromatin structure. For instance, DNA methylation, a critical epigenetic modification, not only regulates gene expression but also plays a key role in the formation of chromatin loops ^19–21^. Chromatin loop prediction algorithms reliant on ChIA-PET data are limited by their dependence on specific target proteins, posing challenges to comprehensively predicting various protein-mediated chromatin interactions ^22–26^.

Nanopore sequencing technology, with its unique advantages in sequencing, has seen increasingly widespread applications in scientific and clinical research ^27–30^. This technology enables the simultaneous detection of DNA sequences and methylation ^31–35^. If a method for predicting chromatin loops based on Nanopore sequencing data could be developed, it would fully leverage the rich information embedded in DNA sequences and epigenetic modifications, thereby achieving more comprehensive and accurate genome-wide chromatin loop predictions.

Such an approach holds the promise of unveiling the intricate regulatory relationships between epigenetic modifications and three-dimensional genome organization, providing new perspectives and deeper insights for related research.

In this study, we present a deep learning framework, NanoLoop, for predicting genome-wide chromatin loops. This framework represents the first attempt to predict the three-dimensional structure of genes using Nanopore sequencing data. By integrating convolutional neural networks and the XGBoost algorithm, NanoLoop identifies key features in DNA sequences and methylation levels, enabling the prediction of heterogeneous chromatin loops regulated by DNA methylation. In tests across four human lymphoblastoid cell lines, NanoLoop demonstrated exceptional performance in chromatin loop prediction and cross-cell-line generalization. Moreover, by fully leveraging methylation information, NanoLoop captures heterogeneous chromatin loops, a capability validated in the regulation of the classic human IGF2-H19 imprinted domain. Further analysis revealed that clustering of methylation at chromatin loop anchor points uncovered four distinct patterns, which are closely associated with histone modifications and chromatin loop types. More importantly, NanoLoop successfully predicted long-range chromatin loops that are challenging for traditional sequence-based methods, accurately identifying 93% of known long-range loops and discovering 44 previously uncharacterized loops. Additionally, NanoLoop revealed methylation-enriched regions (e.g., methylation canyons) and their associations with histone modifications (e.g., H3K27me3). NanoLoop not only provides a convenient tool for Nanopore sequencing users to conduct research from a three-dimensional genomics perspective but also offers robust technical support for large-scale genomic studies and clinical applications. This framework provides new insights into the complex relationship between epigenetic modifications and the three-dimensional genome, holding significant scientific value and practical application potential.

## Results

### The architecture of NanoLoop

NanoLoop utilizes DNA sequence and methylation features derived from Nanopore sequencing data to predict genome-wide chromatin loops. The process of obtaining DNA sequence and methylation involves several steps: raw electrical signals are first base-called to generate sequence signal, which are then error-corrected and assembled into contigs to produce complete DNA sequences. Simultaneously, DNA methylation levels are extracted from the raw signals using deep learning methods. The processed DNA sequences and methylation data are subsequently input into NanoLoop for chromatin loop prediction (**Fig. 1A**).

**Figure 1.**
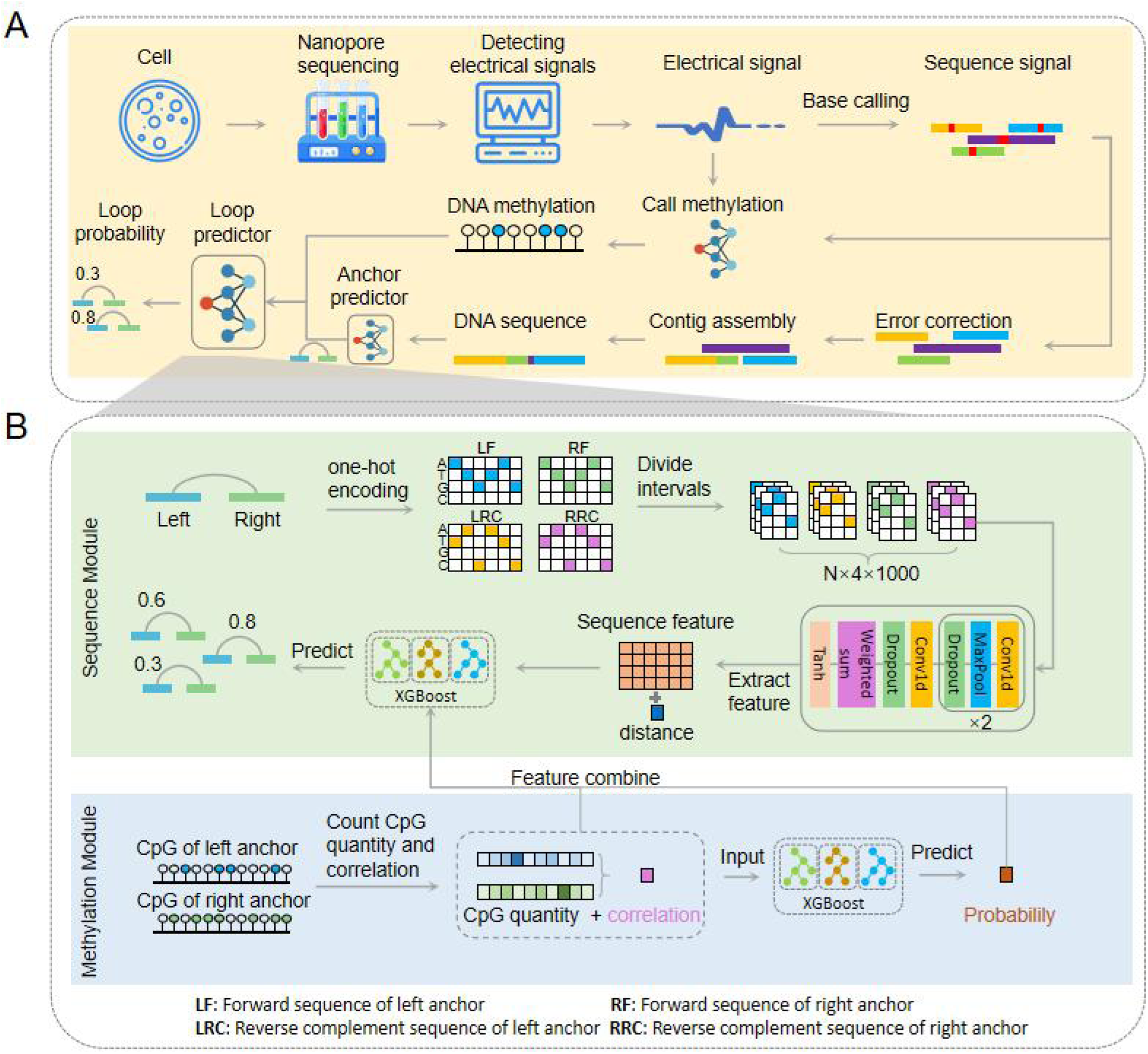
A deep learning framework leveraging Nanopore sequencing for chromatin loop prediction. **(A)** Nanopore sequencing and loop prediction processing. **(B)** NanoLoop integrates DNA sequence and methylation information through a dual-modality architecture.

The core of the NanoLoop architecture comprises the Sequence Module and the Methylation Module, which work in synergy to maximize predictive accuracy (**Fig. 1B**). The Sequence Module processes DNA sequence information from the left and right anchors of potential chromatin loops. Sequences are encoded into a one-hot representation and divided into overlapping intervals, preserving spatial resolution. These intervals are input into a one-dimensional convolutional neural network (Conv1D), which employs ReLU activation and pooling layers to extract sequence-specific features. The resulting 512-dimensional feature vector captures patterns critical to chromatin interactions. This feature vector, along with additional data such as anchor distances, is fed into an XGBoost model to predict loop formation probabilities based on sequence characteristics.

The methylation module employs the XGBoost algorithm to learn the potential relationship between methylation information at chromatin loop anchors and loop formation. Genome-wide analysis reveals that methylation levels at loop anchors show a strong correlation, with most anchors showing methylation levels below 0.6 and a Pearson correlation coefficient of 0.51 between anchor pairs (**Extended Data Figure 1**). Spatially proximal DNA sequences tend to display similar methylation levels, consistent with previous studies ^36–39^. Building on this observation, the module calculates the number of methylated CpG sites within each anchor pair and evaluates the correlation of methylation levels between the left and right anchors. These methylation-based features are then fed into another XGBoost model to generate initial loop predictions. Finally, the methylation features and preliminary prediction probabilities are integrated with the output of the sequence module, and a combined XGBoost model is used to predict the probability of chromatin loop formation. This dual-module architecture effectively captures key features from both DNA sequence and methylation information, enabling comprehensive genome-wide prediction of chromatin loops.

### NanoLoop achieves robust and generalizable chromatin interaction prediction across cell lines

To evaluate NanoLoop’s ability to accurately predict chromatin interactions, we utilized Nanopore sequencing data from the Genome in a Bottle (GIAB) Consortium and the International Genome Sample Resource (IGSR) ^40^. These datasets included four extensively studied cell lines, HG001-HG004, known for their high-quality genomic annotations. The raw electrical signals from Nanopore sequencing were processed through a customized pipeline to generate high-resolution DNA sequences and methylation profiles. Additionally, corresponding Hi-C data for the same samples were used to identify experimentally validated chromatin loops, serving as ground truth labels for model training. This comprehensive integration of DNA sequencing, DNA methylation, and Hi-C provided a robust foundation for NanoLoop’s training process. To ensure that the model’s performance could be evaluated accurately, chromosomes were systematically partitioned into training, testing, and validation sets rather than random partitioning (**Fig. 2A**).

**Figure 2.**
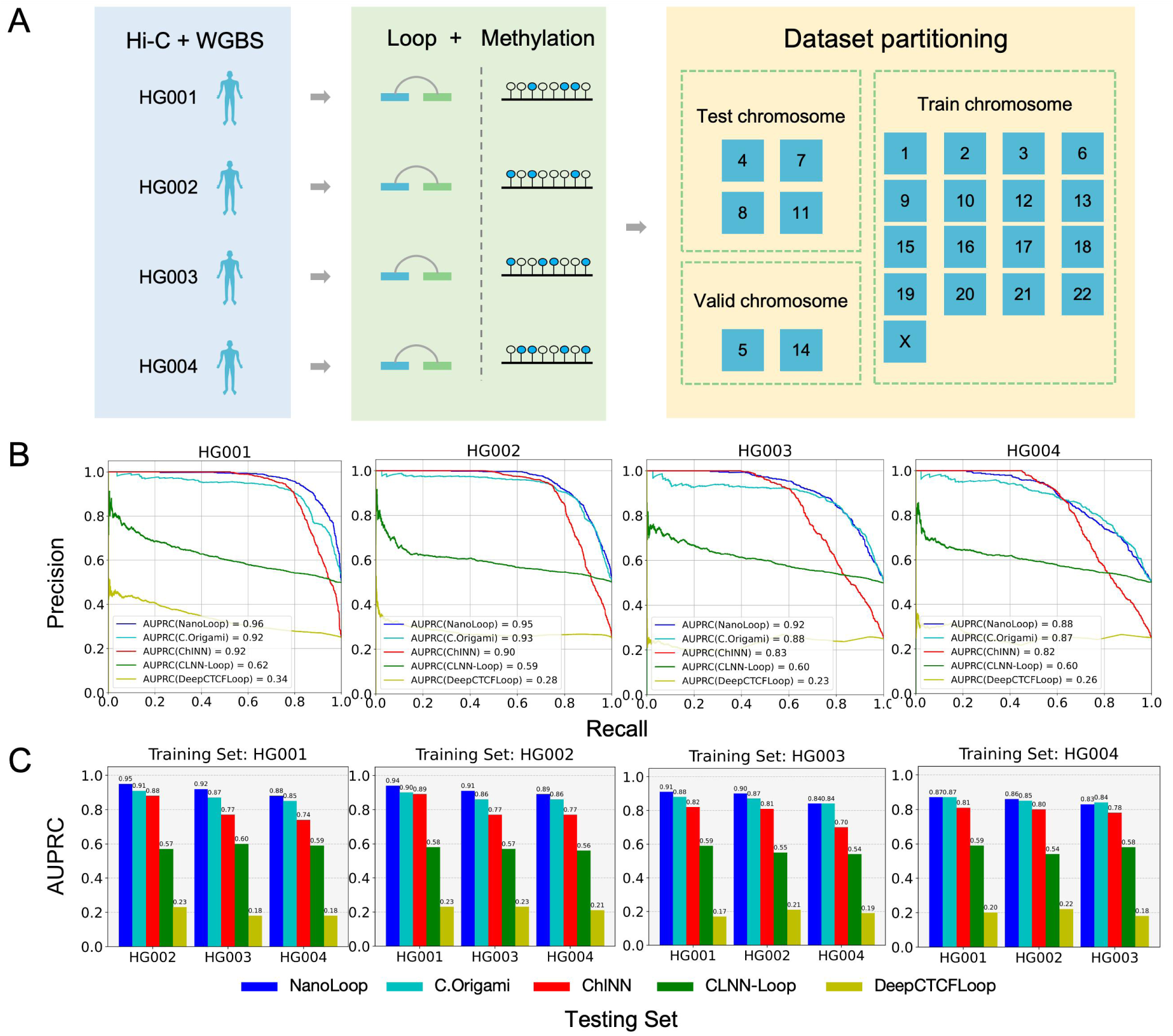
Dataset and Model Performance Comparison with NanoLoop. **(A)** Source of experimental datasets and the chromosome partitioning strategy used for training, validation, and testing. **(B)** Performance of NanoLoop, C.Origami, ChINN, CLNN-Loop and DeepCTCFLoop across cell lines HG001-HG004, evaluated using the Area Under the Precision-Recall Curve (AUPRC). **(C)** Cross-Cell-Line Prediction Performance Comparison of NanoLoop and Other Methods. Models were trained on one cell line and tested on the remaining cell lines.

For benchmarking, we used the same datasets to train existing methods, including C.Origami ^41^, ChINN ^42^, CLNN-Loop^43^, and DeepCTCFLoop^44^, and compared their performance against NanoLoop. Model performance was evaluated using the area under the precision-recall curve (auPRC). NanoLoop consistently outperformed all other methods across all of the four cell lines evaluated, achieving auPRC scores of 0.96, 0.95, 0.92, and 0.88 for HG001, HG002, HG003, and HG004, respectively (**Fig. 2B**).

To assess NanoLoop’s generalization capability, we further evaluated its cross-cell line performance by training the model on one cell line and testing it on the remaining three. NanoLoop demonstrated strong generalization, with auPRC scores exceeding 0.83 across all cross-sample experiments (**Fig. 2C**). In every case, NanoLoop significantly outperformed other models, underscoring its ability to learn robust and transferable loop prediction patterns that generalize effectively to unseen cell lines.

### Distance and methylation correlation drive chromatin loop formation in NanoLoop

To further investigate which features influence chromatin interaction formation, we analyzed the feature importance within NanoLoop. Specifically, we evaluated the model’s performance under four different combinations of sequence, methylation, and distance features (**Fig. 3A**). The results showed that the model achieved the highest auPRC when all features were included, indicating that the integration of multiple feature types is crucial for accurate predictions. In contrast, when the model used only sequence information, its performance was the lowest, with an auPRC of 0.51, suggesting that DNA sequence alone provides limited information for loop prediction. However, the inclusion of methylation and distance features significantly improved the model’s performance, highlighting their substantial influence on loop formation.

**Figure 3.**
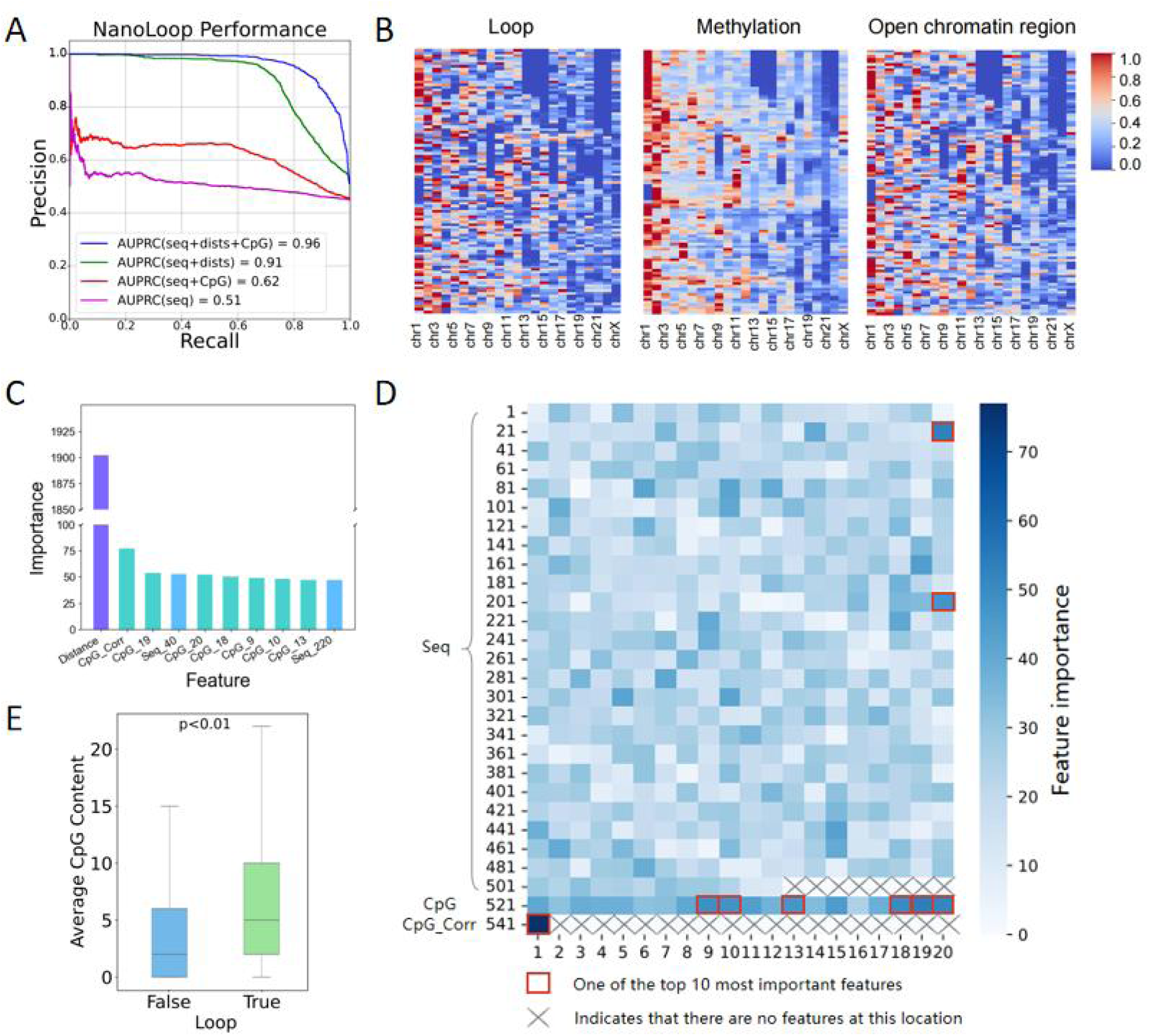
Feature importance analysis in NanoLoop. **(A)** Precision-recall curves of NanoLoop under four feature combinations, integrating sequence, methylation, and distance features. **(B)** Heatmaps showing the genome-wide distribution of 3D chromatin loops, DNA methylation, and open chromatin regions. **(C)** The top 10 most important features identified in the NanoLoop model, including distance, methylation and sequence-related factors. **(D)** Importance heatmap of sequence and methylation features. The top 10 feature importance values are selected using orange boxes, and the blank areas are represented by crosses. **(E)** The average CpG content of positive and negative Loop samples.

To better understand the relationship between chromatin interactions and epigenetic features, we examined the genome-wide distribution of 3D chromatin loops, DNA methylation, and accessible chromatin regions. Each chromosome was divided into 100 equal bins, and the enrichment levels of these features were quantified within these bins (**Fig. 3B**). The analysis revealed that 3D chromatin loops and open chromatin regions exhibited similar clustering patterns across the genome, consistent with previous findings ^45–48^. Furthermore, regions enriched in methylation closely overlapped with loop-enriched regions, suggesting that methylation within proximal regions may play a role in shaping their three-dimensional structure and, in turn, regulating gene expression.

To identify the contribution of individual features to loop prediction, we assessed the importance of all features and listed the top 10 most important features in NanoLoop (**Fig. 3C**). Distance emerged as the most critical feature, with an importance score of 1902, far exceeding other features. Methylation correlation ranked second with an importance score of 77. Among the remaining features, six were methylation-related, and two were sequence-related (**Fig. 3D**). For the non-distance related characteristics, the results revealed that sequence-specific features generally had lower importance, while methylation features were consistently more influential. Overall, methylation correlation was the most significant non-distance feature, indicating that the correlation between anchor pair methylation levels plays a pivotal role in determining loop formation. These findings highlight that, aside from distance, methylation features are the most critical factors for loop prediction. The relatively low importance of sequence features underscores the necessity of incorporating epigenetic information, such as DNA methylation, into sequence-based loop prediction models for enhanced accuracy.

Next, we examined DNA methylation differences between positive and negative samples. The average methylation content at true loop anchors was higher than at randomly generated false loops (**Fig. 3E**), consistent with the idea that chromatin interaction regions tend to accumulate CpG methylation. This observation aligns with prior studies on gene regulation and suggests that DNA methylation plays a role in modulating chromatin accessibility and openness, thus influencing loop formation and gene expression. Moreover, the effect of DNA methylation on chromatin interactions varies by context, as exemplified by the methylation-dependent binding affinities of transcription factors like MeCP2 ^49^. Additionally, we found that exons had the highest median methylation levels among regulatory elements, such as genes, promoters, and enhancers (**Extended Data Figure 2**), indicating that higher methylation in exons may play a role in regulating splicing and RNA processing, while the relatively lower methylation levels in promoters and enhancers may facilitate transcription factor binding, supporting their regulatory functions in gene expression ^50–53^. This highlights the nuanced role of DNA methylation in chromatin structure, where its effects on loop formation are dependent on both the specific genomic region and the context of associated regulatory mechanisms.

### The impact of DNA methylation changes on chromatin loop formation and its application in imprinting domain regulation

DNA methylation dynamics may lead to changes in the three-dimensional structure of the genome, impacting gene expression ^54^. To investigate this, we designed an experiment to test the impact of methylation changes on loop probability. Specifically, we selected equal numbers of loops with predicted probabilities ≥ 0.7 (high-probability group) and ≤ 0.3 (low-probability group). For the low-probability group, we replaced their methylation features with those from the high-probability group while keeping their sequence features unchanged (**Fig. 4A**). We then re-evaluated the loop probabilities for the modified low-probability group (**Fig. 4B**). The results showed a significant increase in loop formation probability after the methylation feature replacement, with some samples reaching probabilities of 0.5 or even 0.8. These findings indicate that changes in DNA methylation levels, independent of sequence alterations, can influence loop formation. This supports the notion that dynamic DNA methylation changes may reshape the three-dimensional genome, enabling dynamic regulation of gene expression in cells. To verify the ability of NanoLoop to predict the impact of DNA methylation changes on loops, we evaluated the regulation of the *IGF2-H19* imprinting domain ^55,56^. *IGF2-H19* imprinting domain regulation is a classic model of DNA methylation affecting chromatin interactions (**Fig. 4C-4D**). The imprinting control regions (ICRs) located approximately 2 kb downstream of *H19* are differentially methylated regions (DMRs). The unmethylated DMRs in the maternal source bind to CTCF, blocking the enhancer(E1)’s regulation of *IGF2* and allowing for the expression of *H19*. However, the DMR methylated in the parent source is unable to bind to CTCF, leading to activation/expression of the *IGF2* gene. After inputting data of DMR unmethylated into NanoLoop, NanoLoop successfully predicted the interaction between the enhancer (E1) and IGF2 on the maternal allele. However, after inputting the data of DMR methylated into NanoLoop, NanoLoop instead predicted the interaction between the enhancer(E2) and *IGF2* on the paternal allele. This indicates that NanoLoop has successfully captured the regulatory relationship between ICR methylation level and loop structure. The patterns learned by NanoLoop may help researchers verify other imprinting domain regulation.

**Figure 4.**
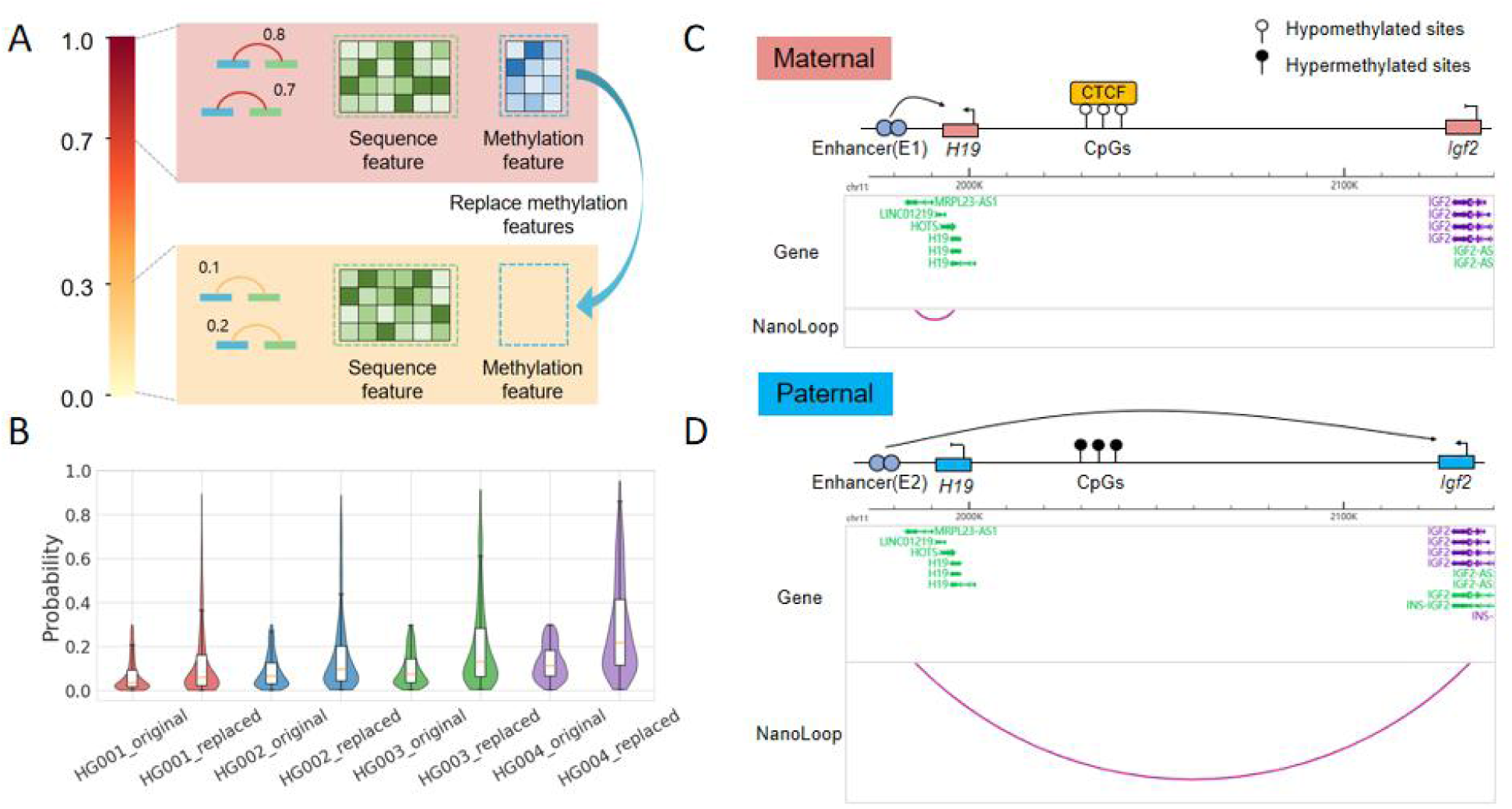
The Effect of DNA Methylation on Loop Formation Probability and an Example of IGF2-H19 Imprinting Domain Regulation. **(A)** Prediction experiment where the methylation features of low-probability loops (≤0.3) were replaced with those from high-probability loops (≥0.7), while sequence features remained unchanged. **(B)** Changes in loop formation probability before and after methylation feature replacement. **(C)** Example of Igf2-H19 imprinting domain regulation based on allelic specific promoter enhancer loop. CTCF binds to unmethylated parent H19 DMR to regulate the expression of H19 by the enhancer. **(D)**. The methylated parent H19 DMR did not bind to CTCF, allowing enhancer to regulate Igf2 expression.

### Clustering analysis reveals the influence of methylation patterns on loop types and TAD structures

To better understand the influence of DNA methylation on loop types, we performed clustering based on the methylation levels of Hi-C loops and anchors in HG001 cells. First, we calculated the Sum of Squared Errors (SSE) for different numbers of clusters to determine the optimal cluster count (**Fig. 5A**). The results indicated that four clusters achieved a good balance between SSE and model complexity. Using the K-Means algorithm, we grouped loops and anchors into four distinct clusters based on their methylation levels (**Fig. 5B-5C**). Although the clustering patterns differed, loops were categorized into upper, lower, left, and right clusters, and anchors were grouped sequentially from left to right. Their overall distributions, however, were quite similar.

**Figure 5:**
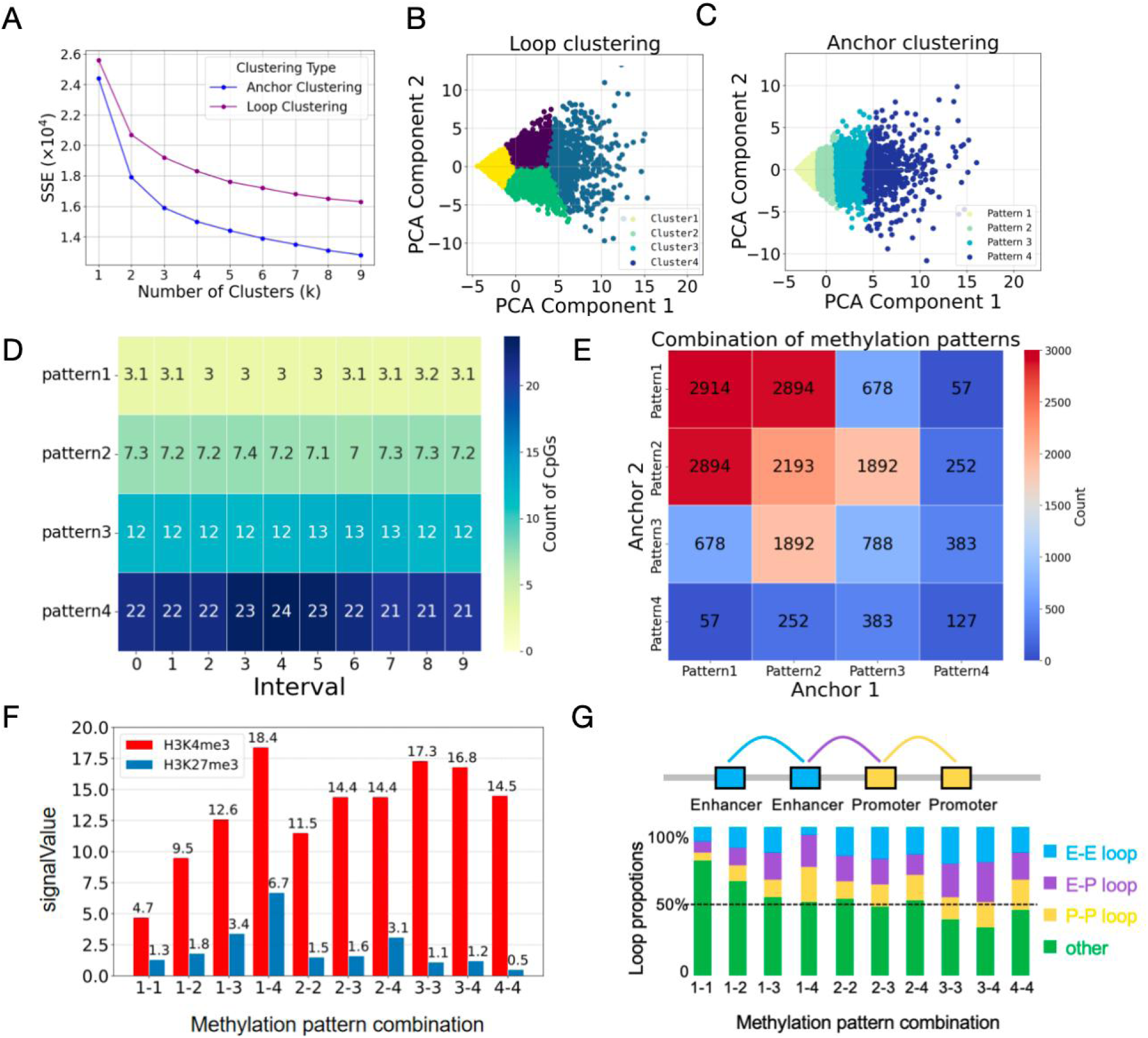
Associations Between Methylation Patterns and Loop Types. **(A)** Elbow plot showing the optimal number of clusters for loop and anchor methylation patterns. **(B)** K-Means clustering of loops based on methylation levels, resulting in four distinct methylation patterns. **(C)** K-Means clustering of anchors based on methylation levels, yielding four different methylation patterns. **(D)** Methylation levels of anchors across the four identified methylation patterns. **(E)** Combinatorial pairing frequencies of the four anchor methylation patterns. **(F)** Enrichment of histone modifications under different anchor methylation pattern combinations. **(G)** Loop types associated with different anchor methylation pattern combinations.

To further investigate DNA methylation patterns, we first divided each anchor into 10 equal bins and calculated the average methylation levels and correlations between bins (**Extended Data Figure 3**). The analysis revealed moderate correlations between bins from different anchors (correlation coefficient ∼0.3), while stronger correlations were observed within the same anchor (correlation coefficient ∼0.6), confirming that spatial proximity within loops contributes to methylation consistency. We then analyzed the average methylation levels across four anchor methylation patterns (**Fig. 5D**). Methylation levels increased progressively from pattern 1 (lowest, with an average of 3 methylation sites per interval) to pattern 4 (highest, with an average of 22 methylation sites per interval). The consistency of methylation levels within each pattern further validated the clustering approach.

To explore potential preferences in methylation pattern combinations, we examined the pairing frequencies of the four anchor methylation clusters (**Fig. 5E**). Most anchor pairs belonged to cluster combinations involving cluster 1, 2, or 3. Notably, cluster 1 anchors preferred pairing with cluster 1 or 2, while cluster 2 anchors frequently paired with clusters 1, 2, or 3. Conversely, anchor pairs with highly divergent methylation patterns (e.g., cluster 1 and cluster 4) were rarely observed, indicating that similar methylation levels between anchors are more conducive to loop formation.

To further elucidate the relationship between methylation combinations and loop formation, we analyzed the enrichment of H3K4me3 and H3K27me3 on loops with different methylation combinations (**Fig.5F**), which are associated with gene activation and silencing, respectively. The results showed significant differences in H3K4me3 enrichment across methylation pattern combinations, with low methylation combinations often having lower H3K4me3 content (such as 1-1, 1-2), while high methylation combinations have relatively higher H3K4me3 content (such as 3-3, 3-4). When the methylation patterns of two anchors differ significantly (such as 1-3, 1-4, 2-4), the enrichment level of H3K4me3 will be higher. When the methylation patterns of the two anchors are close, the enrichment level of H3K4me3 will be relatively lower. Interestingly, in the 1-4 modes with the greatest difference in methylation patterns, H3K4me3 and H3K27me3 have the highest content. This may be due to the high methylation region often recruiting inhibitory histone modifications (H3K27me3), while the low methylation region is easily modified by active histone marks like H3K4me3. Through the coordinated action of DNA methylation writers, readers, and chromatin modifiers, the structural and functional state of chromatin can establish distinct transcriptional regulatory environments between anchor regions. This, in turn, influences the recruitment of histone-modifying enzymes and the enrichment of H3K4me3 and H3K27me3, shaping gene expression outcomes.

We also investigated the loop types associated with different methylation combinations (**Fig. 5G**). Methylation combination cluster 1-1 had the lowest proportions of enhancer-enhancer (E-E), enhancer-promoter (E-P), and promoter-promoter (P-P) loops, totaling only 22.7%. In contrast, higher methylation combinations exhibited increased proportions of these loop types, reaching 68% for cluster 3-4 combinations. These findings suggest that loop methylation levels are strongly associated with regulatory factors like enhancers and promoters, highlighting a potential role for DNA methylation in influencing loop types and regulating gene expression.

### The role of methylation canyons in long-range loop formation and prediction

Studies have shown that large (>7.3 kb), low-DNA-methylation regions, known as methylation canyons, are associated with the formation of long loops, connecting anchors separated by tens of megabases (Mb) ^57^. To investigate the relationship between methylation canyons and long-range loops and explore the influence of DNA methylation on chromatin interactions, we analyzed hematopoietic stem and progenitor cells (HSPCs) ^58,59^. From Hi-C data, we identified 2,683 loops, including 408 manually annotated long-range loops (**Fig. 6A**), and detected 282 methylation canyons using a hidden Markov model. Next, we examined the spatial relationship between long-range loops and methylation canyons (**Fig. 6B**). Among long-range loops, 6.4% have both anchors overlapping with methylation canyons, while 26.5% have one anchor overlapping. The spatial proximity of long-range loops to methylation canyons suggests a potential association between them. We then analyzed the distance distribution of the 408 long-range loops (**Fig. 6C**). The majority (311 loops) spanned less than 2 Mb, with 49 loops spanning 2–4 Mb, and fewer loops at greater distances. Even among long-range loops, the predominant distances remained under 2 Mb. This analysis highlights the close association between long-range loops and methylation canyons.

**Figure 6.**
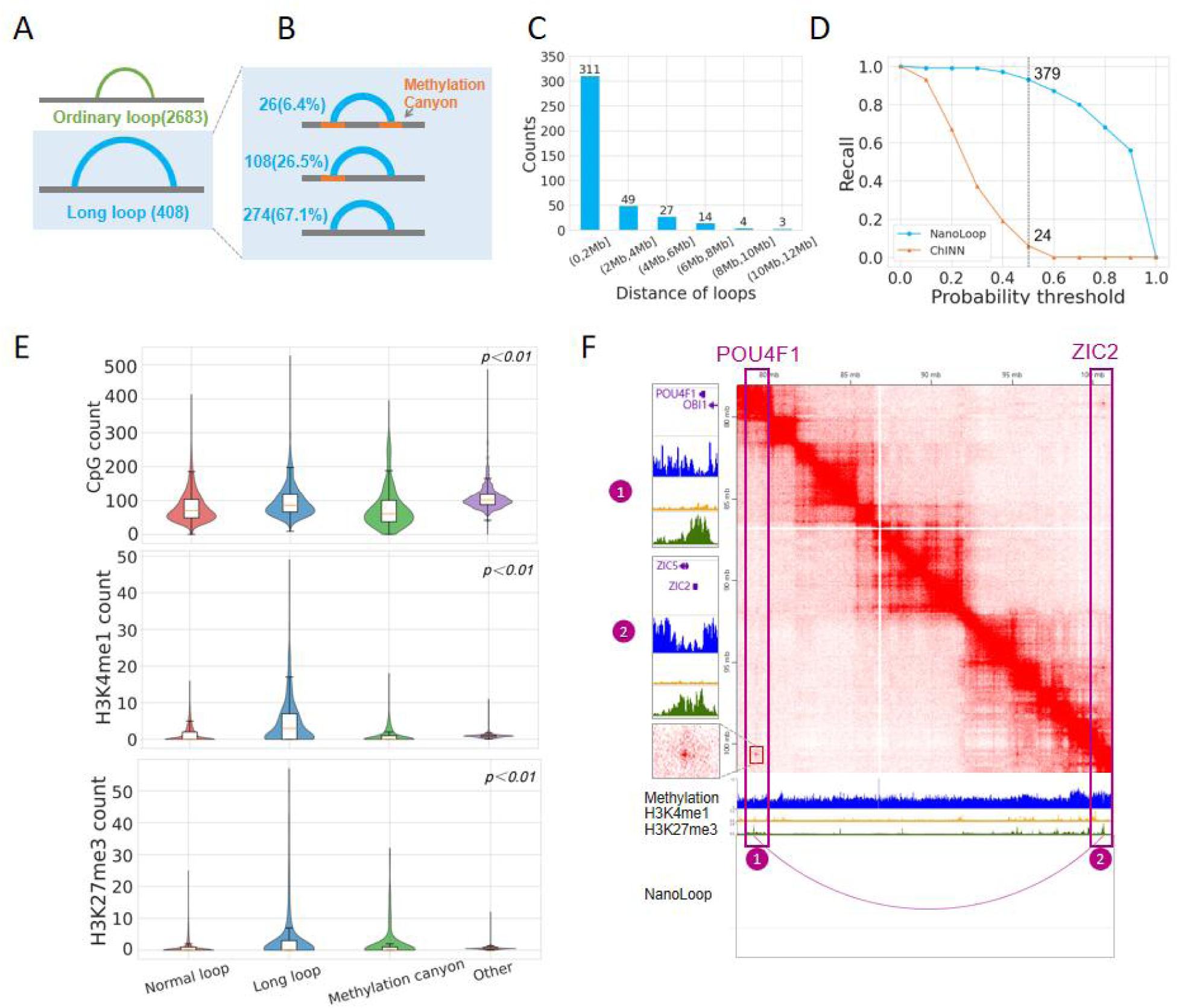
Methylation Canyons and Epigenetic Characteristics of Long-Range Loops in HSPCs. **(A)** Identification of 2683 normal loops and 408 long-range loops from HSPC Hi-C data. **(B)** Three positional relationships between long-range loops and methylation canyons, along with their respective proportions. **(C)** Distance distribution of long-range loops. **(D)** NanoLoop and ChINN predict the recall rate of long-range loops at different probability thresholds. **(E)** Distribution of DNA methylation levels, H3K4me1, and H3K27me3 content across normal loops, long-range loops, methylation canyons, and other regions. **(F)** Long-range interaction between the POU4F1 gene and the ZIC2 gene in HSPCs.

To evaluate NanoLoop’s ability to predict long-range loops, we compared it to ChINN (**Fig. 6D**). We visualized the recall rates of two methods at different probability thresholds. When using a probability of 0.5 as the threshold for distinguishing positive and negative samples, the recall rate of NanoLoop reached 93% (379 samples), while the recall rate of ChINN at the same threshold was only 6% (24 samples). This demonstrates that relying solely on sequence features, as ChINN does, is insufficient for long-range loop prediction. In contrast, NanoLoop integrates DNA methylation features, effectively learning the relationship between methylation canyons and long-range loops, further supporting their strong association.

To compare the epigenomic characteristics of normal loops, long-range loops, methylation canyons, and other regions, we examined the levels of DNA methylation, and H3K4me1 and H3K27me3 enrichment across these four categories (**Fig. 6E**). Long-range loops and methylation canyons exhibited lower methylation levels than normal loops, while normal loops had lower methylation than other regions. This indicates that loop formation generally requires lower methylation levels, and long-range loops demand even lower methylation levels for stability. Additionally, long-range loops showed the highest levels of H3K4me1 and H3K27me3 among the four regions. The high levels of H3K4me1 suggest an open chromatin state, supporting the stability and functionality of long-range loops, enabling physical contacts across genomic regions to establish complex regulatory networks. Conversely, the high levels of the repressive mark, H3K27me3, may help maintain structural stability and prevent unnecessary transcription, and ensure ordered chromatin architecture. These features contribute to the efficient formation and maintenance of long-range loops, which often involve complex regulatory mechanisms such as multiple enhancers targeting the same gene or a single enhancer influencing multiple genes.

As an example of long-range interaction in HSPCs, we identified a loop between *POU4F1* (Chr13:79M) and *ZIC2* (Chr13:100M), two transcription factors involved in developmental processes, particularly neurodevelopment (**Fig. 6F**). Both anchors overlapped with methylation canyons and exhibited strong enrichment of H3K27me3. We have also observed this enrichment phenomenon in many examples of long-range loops (**Extended data Figure 4-6**). NanoLoop successfully predicted this long-range loop, whereas ChINN failed to detect it, further demonstrating NanoLoop’s advantage in incorporating methylation information for accurate prediction of long-range chromatin interactions. We also trained NanoLoop using 408 known long-range loops from HSPCs. To validate the model, we generated a set of random long-range loops across the genome. These random loops were designed to mimic the characteristics of real long-range loops, with similar anchor lengths and loop distances. NanoLoop identified 44 long-range loops from the generated random long-range loops, and upon visual inspection of the visualized Hi-C matrix, it was found that all 44 long-range loops exhibited significant interaction enrichment phenomena(**Supplementary table1**). Therefore, the patterns learned by Nanoloop may help researchers discover new imprinting domains.

## Discussion

Here, we introduce NanoLoop, the first framework to predict genome-wide chromatin loops using nanopore sequencing data, integrating DNA sequence and methylation information through a combination of convolutional neural networks (CNN) and XGBoost. Unlike existing deep learning methods such as C.Origami, ChINN, CLNN-loop, and DeepCTCFLoop, which often depend heavily on sequence-based or CTCF-specific features, NanoLoop harnesses the unique ability of nanopore sequencing to simultaneously capture sequence and methylation signals. This enables a more comprehensive prediction of chromatin interactions, including heterogeneous loops regulated by DNA methylation, a key epigenetic modification known to influence genome folding.

NanoLoop exhibits outstanding predictive performance and cross-cell-line generalization across four human lymphoblastoid cell lines, surpassing sequence-only approaches. By analyzing feature importance, we found that methylation features outweigh sequence-based features, underscoring the necessity of incorporating epigenetic data for accurate loop prediction. Leveraging the epigenetic information provided by methylation, NanoLoop successfully predicts heterogeneous loops, a capability validated in the regulation of the IGF2-H19 imprinted domain. Notably, NanoLoop identifies four distinct methylation patterns at loop anchors, which correlate with histone modifications (e.g., H3K4me3 and H3K27me3) and loop types (e.g., enhancer-promoter interactions), revealing how methylation shapes chromatin architecture beyond CTCF-mediated boundaries.

A standout feature of NanoLoop is its ability to predict long-range chromatin loops, a challenge for traditional methods. It successfully identified 93% of known long-range loops in HSPCs and uncovered 44 previously uncharacterized loops, validated through Hi-C heatmap analysis. We also observed associations between long-range loops and methylation canyons, with lower methylation levels and H3K27me3 enrichment suggesting a cooperative role in loop formation. These findings highlight NanoLoop’s potential to uncover novel regulatory mechanisms inaccessible to sequence-only models.

In the future, as nanopore sequencing technology continues to evolve, its ability to capture increasingly precise DNA sequence and epigenetic information will enable the discovery of a broader range of biological insights. With the steady accumulation of sequencing samples, comprehensive analyses of vast datasets will deepen our understanding of the intricate interplay between DNA methylation and chromatin three-dimensional structure. Furthermore, by integrating an expanding collection of biological experimental data, such as from cell lines and animal models, alongside clinical samples, NanoLoop is poised to play a pivotal role in unraveling the mechanisms of disease. NanoLoop will drive advancements in fields such as three-dimensional genomics, epigenetic regulatory network construction, and the identification of disease biomarkers, ultimately providing critical support for the development of precision medicine.

## Materials and methods

### Nanopore processing flow

Nanopore Sequencing Data Processing Pipeline. In this study, we developed a comprehensive pipeline for processing Oxford Nanopore Technologies (ONT) sequencing data, with the goal of predicting anchor points in DNA sequences for subsequent loop prediction. The pipeline consists of four main stages: base calling, assembly, methylation calling, and anchor prediction. Each stage is designed to progressively refine the raw sequencing data into high-quality, informative outputs that can be used for downstream analysis.

1. Base calling: The fast5 file obtained from the raw sequencing of Nanopore is an electrical signal that needs to be converted into bases through base calling. The guppy basecaller method in the guppy toolkit can be used to perform base calls on the Oxford Nanopore Technologies sequencing data.
2. Nanopore assembly: The fastq file after basecalling needs to be assembled to generate longer DNA sequences for subsequent loop prediction. The commonly used assembly tool for Nanopore is Flye ^60^, Flye is a de novo assembler used for single molecule sequencing reading, which can read raw ONTs as input and output optimized overlapping populations. The brief assembly command is:

Flye --nano-raw HG003.fastq.gz --out-dir /path/to/output_directory --threads 40 --genome-size 3g.

1. DNA methylation information can be identified from Nanopore’s sequence and electrical signals, with Deepsignal and Nanopolish being classic DNA methylation recognition methods. Deepsignal, in particular, is a deep learning-based approach used to detect DNA methylation states from Oxford Nanopore sequencing reads. After basecalling with Guppy and signal reshuffling with Tombo, Nanopore sequencing signals can be input into Deepsignal for methylation recognition.
2. Predicting anchors: In our previous work, we proposed a method for predicting anchors that can predict potential anchor sites from input DNA sequences. Specifically, the input DNA sequence is processed through convolution blocks and Transformer blocks to extract features.

Four different convolution operations are performed to generate four orbital signals, and the significant peaks of the orbital signals are identified through peak calling, thereby determining potential anchors and providing anchor site data for downstream loop prediction.

### Nanoporeloop prediction

NanoLoop consists of a methylation module and a sequence module. The methylation module initially predicts loop probability from methylation features, while the sequence module extracts features from DNA sequences and uses methylation, distance, and sequence features for final prediction.

1. The Methylation module is composed of XGBoost (Extreme Gradient Boosting), which is a machine learning algorithm based on Gradient Boosting Decision Trees (GBDT). By combining multiple weak learners, it can capture complex nonlinear relationships and provide higher prediction accuracy. The input data for the Methylation module consists of 20 dimensional CpG features and 1-dimensional methylation correlation. Based on the methylation information, the Methylation module will preliminarily predict the probability of anchor pairs forming loops. The model input is then concatenated with the output probability to form a 22-dimensional feature, denoted as Output 1, which serves as input for the Sequence module.
2. The sequence module consists of a sequence feature extractor composed of CNN (Convolutional Neural Network) and a regressor composed of XGBoost. The beginning of the sequence feature extractor has two feature convolution blocks, which have Conv1d, maxpool, and dropout. Conv1d is responsible for scanning and extracting local patterns on DNA sequences, maxpool can perform max pooling on feature maps to reduce their spatial dimensions, and dropout can randomly discard a portion of neurons during training to improve the model’s generalization ability. Two feature convolution blocks are followed by one Conv1d and dropout, as well as weighted sum and Tanh activation functions. Weighted_Sum is a learnable parameter matrix that can compress the features extracted by convolutional blocks. In backpropagation, the model can automatically adjust these weights to maximize the contribution of each convolutional channel to different targets. The weighted sum result is processed by the Tanh activation function to obtain a 128-dimensional Output 2. After concatenating Output1 and Output2, they will be input into the XGBoost regressor for prediction. The regressor will output a probability value between 0-1, representing the probability of anchor pairs forming a loop.

### NanoLoop Training Workflow

The training of the NanoLoop model is carried out in three stages. In the first stage, the methylation module is trained using the 20 dimensional methylation features of anchors and the 1-dimensional methylation correlation, with the goal of outputting the probability of anchor pairs forming loops. The input and output of the methylation module will be concatenated into 22 dimensional features for prediction in the third stage. In the second stage, the positive DNA sequences of the left and right anchors will be input into the sequence module. The sequence module will first generate reverse sequences based on the forward DNA sequence, resulting in four sets of sequences. Each sequence will be processed into a 4-dimensional matrix through one hot encoding, and then the matrix will be divided into several 1000bp intervals, with adjacent intervals overlapping by 500 bp. Each set of sequences will be fed into a feature extractor for training, resulting in 128 dimensional features. A total of 512 dimensional sequence features will be obtained from 4 sets of sequences. These features will be input into a fully connected layer to predict the probability of loop formation. The parameters of the feature extractor will be trained during backpropagation, and after training, the parameters of the feature extractor will be frozen. In the third stage, the fully connected layer is replaced by the XGBoost model, and the 22 dimensional features of the methylation module, the 512 dimensional sequence features output by the sequence feature extractor, and the 1-dimensional distance features are connected into a 535 dimensional feature vector. This 535-dimensional feature vector is then input into the XGBoost regressor to predict the probability of loop formation, providing a robust framework for identifying chromatin interactions.

### Methylation feature construction

Methylation sites are sparsely distributed across the genome. To address this, we evenly divided each anchor into 10 intervals and counted the number of CpG sites with a methylation probability of ≥ 0.5 in each interval. That is, one anchor pair can generate two 10 dimensional methylation features. Next, calculate the Pearson correlation coefficient for the 10 dimensional features (X) of the left anchor and the 10 dimensional features (Y) of the right anchor. The formula is:

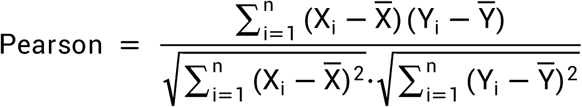

X_i_ and Y_i_ are the i-th observations of variables X and Y, X and Y are the means of variables X and Y,respectively. n is the number of observations, which is 10 in this experiment.

### Positive and negative sample generation strategy

(1) Positive sample generation strategy. Firstly, use the Juicer tool to convert the raw fastq data obtained from the Hi-C experiment into. hic data. Then, generate the corresponding restriction enzyme file using generate_site_position.py, and identify the chromatin ring from the. hic data using the HiCCUPS tool ^61^ in juicer tools. The key parameters include: java -Xmx20g
-jar juicer.tools.jar hiccups - threads 40.
(2) Negative sample generation strategy.Because most of the loops called by Hiccups have a resolution of 5k, we generate negative anchors by sliding a 5000bp window on the genome and ensure that there is at least a 500 bp gap between the negative anchor and the positive anchor. The generated negative anchors are randomly paired to generate negative loops, and the negative loops are sampled according to the distribution ratio of the positive loops at different distances, so that the distance distribution of the negative loops is as close as possible to the positive loops, ensuring that the final number of generated negative and positive samples is close to 1:1 balance.

### Loop Prediction Using C.Origami

We performed loop calling analysis on C.Origami-predicted Hi-C contact matrices at 10-kb resolution with interaction distance parameters set between 30 kb and 1 Mb. The predicted matrices were converted to valid pairs by merging chromosome-wise predictions and calculating discretized interaction intensity values, which were then processed using FitHiC to identify significant chromatin interactions with FDR-adjusted Q-values. To evaluate prediction accuracy, we filtered significant loops using Q-value cutoffs ranging from 1×10^-5^ to 1×10^-13^ and compared them with experimentally derived loops from the reference dataset to calculate AUPRC.

### Loop Prediction Using ChINN

ChINN’s training data is consistent with NanoLoop. In terms of loop partitioning, chromosomes 4, 7, 8, and 11 are used as training sets, chromosomes 5 and 14 are used as validation sets, and the remaining chromosomes are used as test sets. We also use the third-generation sequencing data we assembled as DNA sequence data. The input of the model is the DNA sequence of each anchor pair. We trained the feature extractor using train_distance_matched.py and the classifier using train_extended.py, with default parameters, and then made predictions on the validation set. The final predicted result is the probability of each anchor pair forming a loop. We use 0.5 as the threshold to distinguish between positive and negative samples.

### Loop Prediction Using DeepCTCFLoop

Input data: The input data is a 1038bp DNA sequence stored in FASTA format. We selected loops containing CTCF binding sites from the positive samples of NanoLoop and extracted 1038bp of anchor centers as positive samples. Then, a 1038bp sequence was extracted from the anchor center of the negative sample in NanoLoop as the negative sample.

Model training: Convert the DNA sequence into a heat encoded 4D matrix and input it into the model. Set the parameters to a learning rate of 0.0001, a batch size of 16, and a training epoch of 100. The training process is handled by the evaluation() function in train.exe. The model undergoes multiple rounds of training to ensure robust generalization. If the validation loss does not improve within 5 iteration cycles, please stop training as soon as possible to avoid overfitting. During the training process, the best model weights will be saved when the validation loss is at the lowest point.

### Loop Prediction Using CLNN-loop

Input data: The input data is a 42bp DNA sequence stored in FASTA format. We selected loops containing CTCF binding sites from the positive samples of Nanoloops and extracted 42bp of anchor centers as positive samples. Next, a 42bp sequence was extracted from the anchor center of the negative sample in NanoLoop as the negative sample.

Model training: First, input the fasta file into feature_code.py to extract features, and then use data-load.py to fuse the extracted multiple features to generate the final input feature matrix. The core parameters for training LSTM are set as follows: learning rate: 0.001, batch size: 50, epoch count: 150, and other parameters follow the default settings. When the model’s performance stops improving, training is halted, and ModelCheckpoint is used to save the best model.

### Clustering of loops

(1)We used the 20-dimensional CpG features of each loop, derived from the left and right anchors as described in the “Methylation feature construction” section, as the input features for clustering. These features were normalized to have zero mean and unit variance before being input into the K-means algorithm, with the number of clusters (’n_clusters’) set to 4.

The standardized formula is:

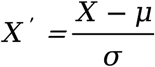

Where μ is the sample mean and σ is the sample standard deviation.

(2) Clustering of anchors: We use the 10 dimensional CpG features of anchors as a sample, meaning that one loop can generate two samples. Then standardize the zero mean and unit variance of the 10 dimensional CpG feature features, and input them into the K-means method for clustering, with a specified value of 4 for the clusters.

### Identification of methylation canyon

The long-range and short-range loops in HSPC, as well as histone modification data, are from the appendix data of Zhang et al ^59^. The short-range loop is annotated with default Juicer parameters using HICCUPS at resolutions of 5kB and 10kB for standard loops. Long range loops are selected by visually inspecting loops with a resolution of 25-100kb using a balanced normalization mode, excluding loops within 2 Mb of the diagonal.

## Extended data

**Extended Data Fig. 1.**
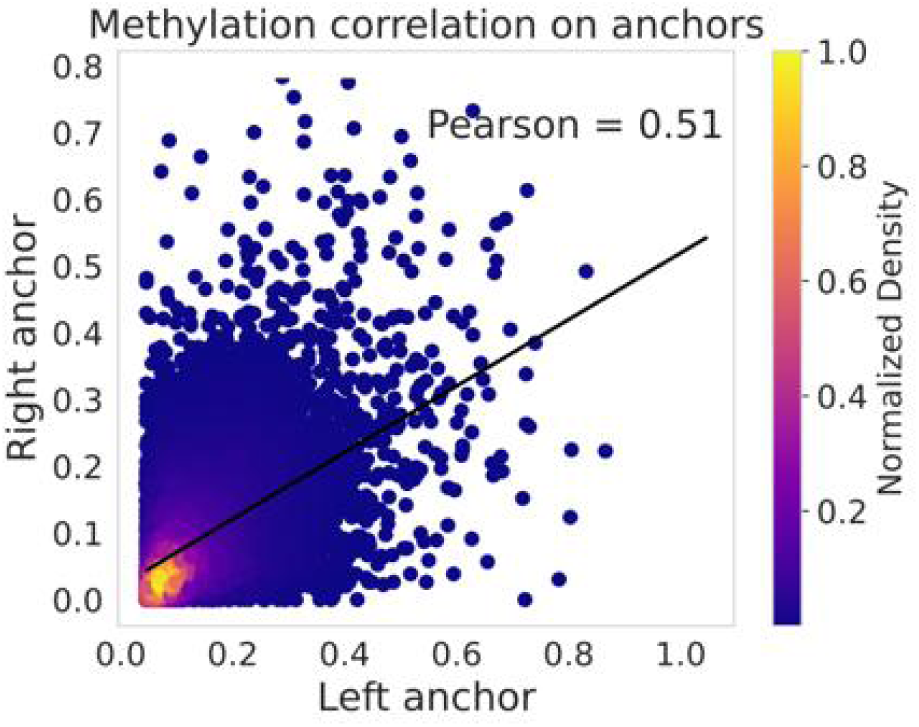
Distribution of DNA methylation levels across anchor pairs. Each point represents a loop, with the x- and y-axes indicating the methylation levels of the left and right anchors, respectively. The overall Pearson correlation coefficient for anchor pair methylation levels is 0.51.

**Extended Data Fig. 2.**
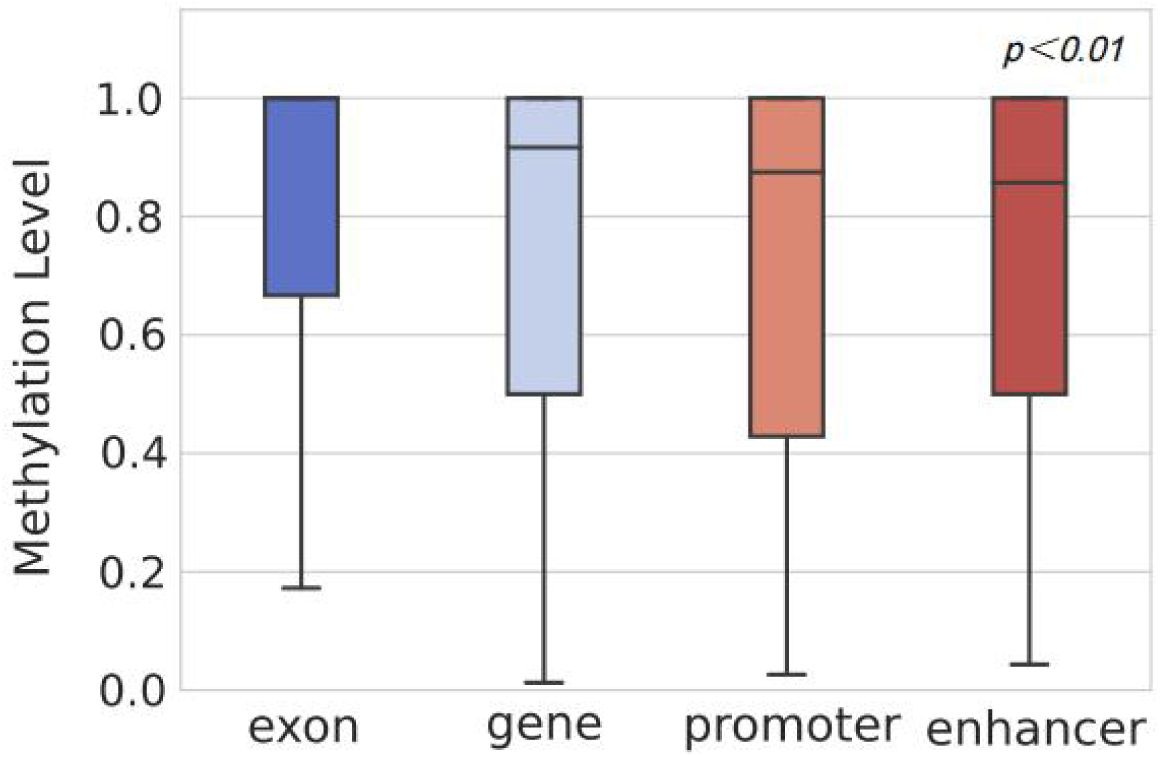
methylation status of regulatory elements. The methylation probability of CpG sites within exon, gene, promoter, and enhancer regions was calculated as the methylation level.

**Extended Data Fig. 3.**
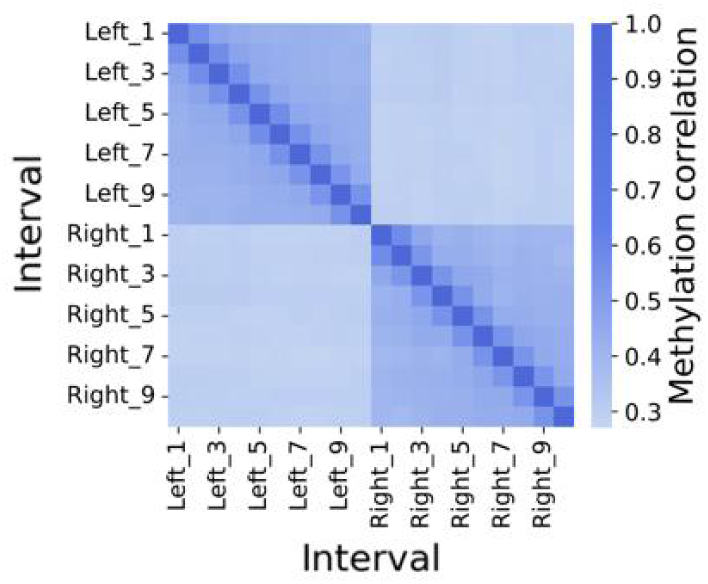
Correlation of average methylation levels across different bins of anchor pairs. Left_1 to Left_10 represent the left anchor evenly divided into 10 intervals, and Right_1 to Right_10 represent the right anchor similarly. The methylation level is calculated as the proportion of methylation sites with a frequency ≥0.5 in each interval, and the methylation correlation between intervals is then computed.

**Extended Data Fig. 4.**
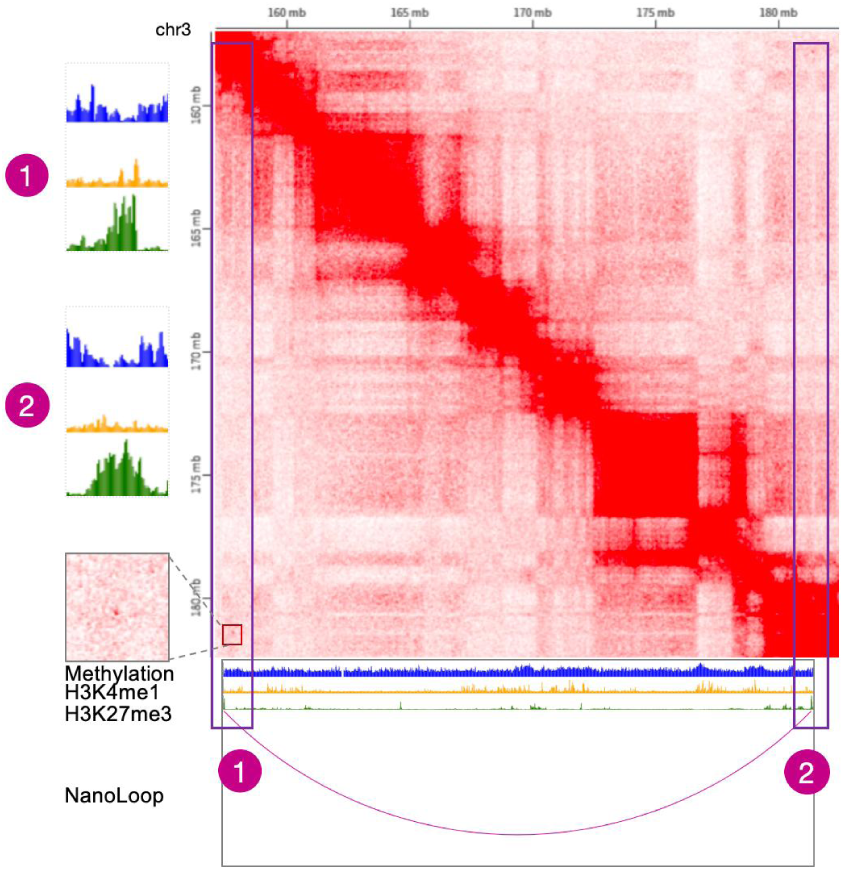
NanoLoop-Predicted Long-Range Chromatin Loops on Chromosome 3 with Epigenetic Signal Enrichment. Hi-C heatmap showing a NanoLoop-predicted long-range chromatin loop on chromosome 3 in HSPCs, spanning about 25 Mb. Tracks 1 and 2 display reduced methylation and H3K27me3 enrichment at loop anchors.

**Extended Data Fig. 5.**
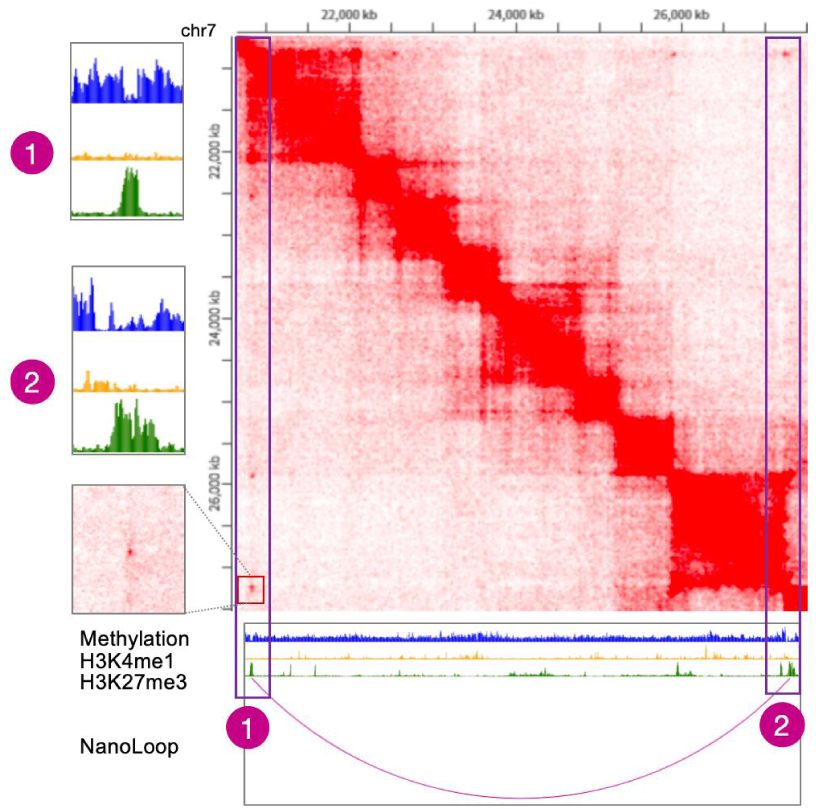
NanoLoop-Predicted Long-Range Chromatin Loops on Chromosome 7 with Epigenetic Signal Enrichment. Hi-C heatmap showing a NanoLoop-predicted long-range chromatin loop on chromosome 7 in HSPCs, spanning about 6 Mb. Tracks 1 and 2 display reduced methylation and H3K27me3 enrichment at loop anchors.

**Extended Data Fig. 6.**
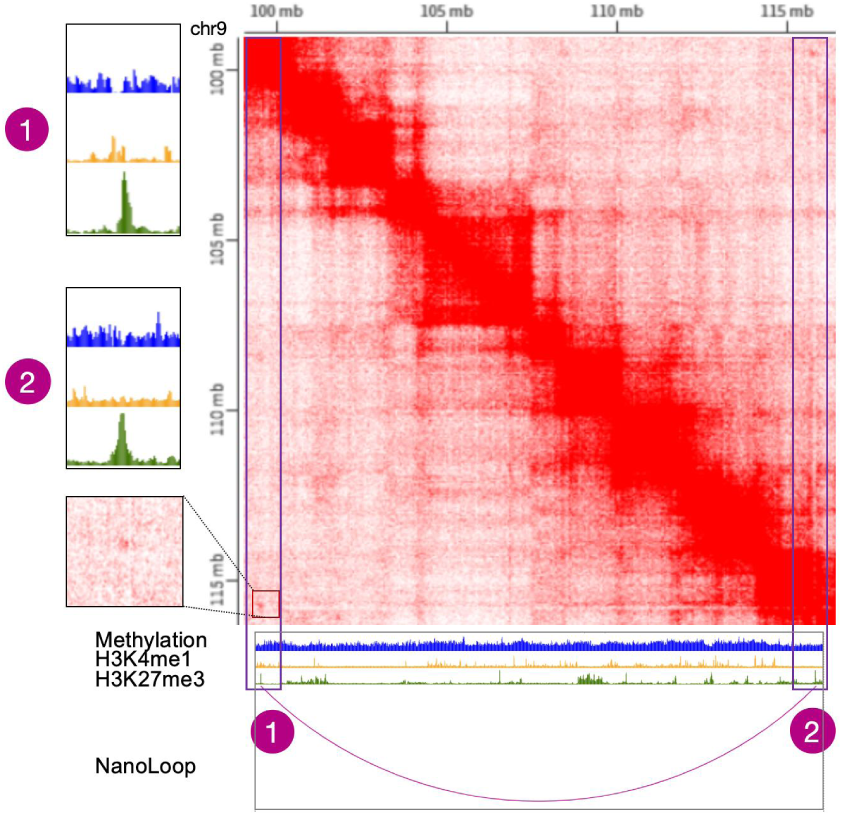
NanoLoop-Predicted Long-Range Chromatin Loops on Chromosome 9 with Epigenetic Signal Enrichment. Hi-C heatmap showing a NanoLoop-predicted long-range chromatin loop on chromosome 9 in HSPCs, spanning about 15 Mb. Tracks 1 and 2 display reduced methylation and H3K27me3 enrichment at loop anchors.

## Supplementary Materials

Supplementary table 1. NanoLoop predicts long-range loops from HSPCs.xlsx

## Acknowledgements

This work was supported by grants from the National Natural Science Foundation of China under Grants (No. 62225209, No. 62320106009 to M.L.). We are grateful to the High-Performance Computing Center of Central South University for partial support of this work.

## Author contributions

W.H., L.T. and M.L. conceived of the presented idea. W.H. collected the data and designed the model of NanoLoop. W.H., L.T. and M.C.H. helped improve the bioinformatics analysis. L.T., M.C.H. and M.L. aided in interpreting the results and provided input on the data presentation. Y. Z assisted in testing the comparison method, while J.X helped draw charts and checked for the presence of long-range loops. All authors provided critical feedback and helped shape the research, analysis and manuscript.

## References

1. Zheng, M. et al. Multiplex chromatin interactions with single-molecule precision. Nature 566, 558–562 (2019).

2. Li, G. et al. Extensive promoter-centered chromatin interactions provide a topological basis for transcription regulation. Cell 148, 84–98 (2012).

3. Tang, Z. et al. CTCF-Mediated Human 3D Genome Architecture Reveals Chromatin Topology for Transcription. Cell 163, 1611–1627 (2015).

4. Weintraub, A. S. et al. YY1 Is a Structural Regulator of Enhancer-Promoter Loops. Cell 171, 1573–1588.e28 (2017).

5. Bonev, B. & Cavalli, G. Organization and function of the 3D genome. Nat Rev Genet 17, 661–678 (2016).

6. Fraser J, Williamson I, Bickmore WA, Dostie J. An overview of genome organization and how we got there: From FISH to Hi-C. Microbiol Mol Biol Rev. 2015;79:347 – 372.

7. Spitz F. Gene regulation at a distance: From remote enhancers to 3D regulatory ensembles. Semin Cell Dev Biol. 2016;57:57 – 67.

8. Lieberman-Aiden, E. et al. Comprehensive Mapping of Long-Range Interactions Reveals Folding Principles of the Human Genome. Science 326, 289–293 (2009).

9. Fullwood, M. J. et al. An oestrogen-receptor-α-bound human chromatin interactome. Nature 462, 58 (2009).

10. Fang, R. et al. Mapping of long-range chromatin interactions by proximity ligation-assisted ChIP-seq. Cell Res 26, 1345 (2016).

11. Mumbach, M. R. et al. HiChIP: efficient and sensitive analysis of protein-directed genome architecture. Nat Methods 13, 919–922 (2016).

12. Dsouza K B, Maslova A, Al-Jibury E, et al. Learning representations of chromatin contacts using a recurrent neural network identifies genomic drivers of conformation[J]. Nature Communications, 2022, 13(1): 3704.

13. Lin D, Hong P, Zhang S, et al. Digestion-ligation-only Hi-C is an efficient and cost-effective method for chromosome conformation capture[J]. Nature genetics, 2018, 50(5): 754–763.

14. Díaz N, Kruse K, Erdmann T, et al. Chromatin conformation analysis of primary patient tissue using a low input Hi-C method[J]. Nature communications, 2018, 9(1): 4938.

15. Salameh T J, Wang X, Song F, et al. A supervised learning framework for chromatin loop detection in genome-wide contact maps[J]. Nature communications, 2020, 11(1): 3428.

16. Yu M, Abnousi A, Zhang Y, et al. SnapHiC: a computational pipeline to identify chromatin loops from single-cell Hi-C data[J]. Nature methods, 2021, 18(9): 1056–1059.

17. Wolff J, Backofen R, Grüning B. Loop detection using Hi-C data with HiCExplorer[J]. Gigascience, 2022, 11: giac061.

18. Zhang S, Plummer D, Lu L, et al. DeepLoop robustly maps chromatin interactions from sparse allele-resolved or single-cell Hi-C data at kilobase resolution[J]. Nature genetics, 2022, 54(7): 1013–1025.

19. Monteagudo-Sánchez A, Noordermeer D, Greenberg M V C. The impact of DNA methylation on CTCF-mediated 3D genome organization[J]. Nature Structural & Molecular Biology, 2024, 31(3): 404–412.

20. Buitrago D, Labrador M, Arcon J P, et al. Impact of DNA methylation on 3D genome structure[J]. Nature Communications, 2021, 12(1): 3243.

21. Lee D S, Luo C, Zhou J, et al. Simultaneous profiling of 3D genome structure and DNA methylation in single human cells[J]. Nature methods, 2019, 16(10): 999–1006.

22. Liu T, Wang Z. DeepChIA-PET: Accurately predicting ChIA-PET from Hi-C and ChIP-seq with deep dilated networks[J]. PLOS Computational Biology, 2023, 19(7): e1011307.

23. Kai Y, Andricovich J, Zeng Z, et al. Predicting CTCF-mediated chromatin interactions by integrating genomic and epigenomic features[J]. Nature communications, 2018, 9(1): 4221.

24. Cao Q, Anyansi C, Hu X, et al. Reconstruction of enhancer – target networks in 935 samples of human primary cells, tissues and cell lines[J]. Nature genetics, 2017, 49(10): 1428–1436.

25. He B, Chen C, Teng L, et al. Global view of enhancer – promoter interactome in human cells[J]. Proceedings of the National Academy of Sciences, 2014, 111(21): E2191–E2199.

26. Whalen S, Truty R M, Pollard K S. Enhancer – promoter interactions are encoded by complex genomic signatures on looping chromatin[J]. Nature genetics, 2016, 48(5): 488–496.

27. Xiao, Tiantian, and Wenhao Zhou. “The third generation sequencing: the advanced approach to genetic diseases.” Translational pediatrics 9.2 (2020): 163.

28. Gilpatrick, Timothy, et al. “Targeted nanopore sequencing with Cas9-guided adapter ligation.” Nature biotechnology 38.4 (2020): 433–438.

29. Lin, Bo, Jianan Hui, and Hongju Mao. “Nanopore technology and its applications in gene sequencing.” Biosensors 11.7 (2021): 214.

30. Kono, Nobuaki, and Kazuharu Arakawa. “Nanopore sequencing: Review of potential applications in functional genomics.” Development, growth & differentiation 61.5 (2019): 316–326.

31. Ni, P. et al. DeepSignal: detecting DNA methylation state from Nanopore sequencing reads using deep-learning. Bioinformatics 35, 4586–4595 (2019).

32. Jain, M., Olsen, H. E., Paten, B. & Akeson, M. The Oxford Nanopore MinION: delivery of nanopore sequencing to the genomics community. Genome Biol. 17, 239 (2016).

33. Rand, A. C. et al. Mapping DNA Methylation with High Throughput Nanopore Sequencing. Nat. methods 14, 411–413 (2017).

34. Stoiber, M., et al. De novo Identification of DNA Modifications Enabled by Genome-Guided Nanopore Signal Processing. bioRxiv 094672 (2017) doi:10.1101/094672.

35. Liu, Q., Georgieva, D. C., Egli, D. & Wang, K. NanoMod: a computational tool to detect DNA modifications using Nanopore long-read sequencing data. BMC Genom. 20, 78 (2019).

36. Li G, Liu Y, Zhang Y, et al. Joint profiling of DNA methylation and chromatin architecture in single cells[J]. Nature methods, 2019, 16(10): 991–993.

37. Guo S, Diep D, Plongthongkum N, et al. Identification of methylation haplotype blocks aids in deconvolution of heterogeneous tissue samples and tumor tissue-of-origin mapping from plasma DNA[J]. Nature genetics, 2017, 49(4): 635–642.

38. Shoemaker R, Deng J, Wang W, et al. Allele-specific methylation is prevalent and is contributed by CpG-SNPs in the human genome[J]. Genome research, 2010, 20(7): 883–889.

39. Dixon J R, Gorkin D U, Ren B. Chromatin domains: the unit of chromosome organization[J]. Molecular cell, 2016, 62(5): 668–680.

40. Fairley S, Lowy-Gallego E, Perry E, et al. The International Genome Sample Resource (IGSR) collection of open human genomic variation resources[J]. Nucleic acids research, 2020, 48(D1): D941–D947.

41. Tan J, Shenker-Tauris N, Rodriguez-Hernaez J, et al. Cell-type-specific prediction of 3D chromatin organization enables high-throughput in silico genetic screening[J]. Nature biotechnology, 2023, 41(8): 1140–1150.

42. Cao F, Zhang Y, Cai Y, et al. Chromatin interaction neural network (ChINN): a machine learning-based method for predicting chromatin interactions from DNA sequences[J]. Genome biology, 2021, 22: 1–25.

43. Zhang P, Wu Y, Zhou H, et al. CLNN-loop: a deep learning model to predict CTCF-mediated chromatin loops in the different cell lines and CTCF-binding sites (CBS) pair types[J]. Bioinformatics, 2022, 38(19): 4497–4504.

44. Kuang S, Wang L. Deep learning of sequence patterns for CCCTC-binding factor-mediated chromatin loop formation[J]. Journal of Computational Biology, 2021, 28(2): 133–145.

45. Hsiung C C S, Wilson C M, Sambold N A, et al. Engineered CRISPR-Cas12a for higher-order combinatorial chromatin perturbations[J]. Nature Biotechnology, 2024: 1–15.

46. Zhang Y, Dong Q, Wang Z, et al. A fine-scale Arabidopsis chromatin landscape reveals chromatin conformation-associated transcriptional dynamics[J]. Nature Communications, 2024, 15(1): 3253.

47. Chiariello A M, Abraham A, Bianco S, et al. Multiscale modelling of chromatin 4D organization in SARS-CoV-2 infected cells[J]. Nature Communications, 2024, 15(1): 4014.

48. Daugird T A, Shi Y, Holland K L, et al. Correlative single molecule lattice light sheet imaging reveals the dynamic relationship between nucleosomes and the local chromatin environment[J]. Nature Communications, 2024, 15(1): 4178.

49. Bajikar S S, Zhou J, O’Hara R, et al. Acute MeCP2 loss in adult mice reveals transcriptional and chromatin changes that precede neurological dysfunction and inform pathogenesis[J]. Neuron, 2025, 113(3): 380–395. e8.

50. Bird, A. P. & Wolffe, A. P. Methylation-induced repression–belts, braces, and chromatin. Cell 99, 451–454 (1999).

51. Siegfried, Z. et al. DNA methylation represses transcription in vivo. Nat. Genet. 22, 203–206 (1999).

52. Kulis, M., Queiros, A. C., Beekman, R. & Martin-Subero, J. I. Intragenic DNA methylation in transcriptional regulation, normal differentiation and cancer. Biochim Biophys. Acta 1829, 1161–1174 (2013).

53. Suzuki, M. M. & Bird, A. DNA methylation landscapes: provocative insights from epigenomics. Nat. Rev. Genet. 9, 465–476 (2008).

54. Liu H, Zhou J, Tian W, et al. DNA methylation atlas of the mouse brain at single-cell resolution[J]. Nature, 2021, 598(7879): 120–128.

55. Matsuzaki H, Miyajima Y, Fukamizu A, et al. Orientation of mouse H19 ICR affects imprinted H19 gene expression through promoter methylation-dependent and-independent mechanisms[J]. Communications Biology, 2021, 4(1): 1410.

56. Venkatraman A, He X C, Thorvaldsen J L, et al. Maternal imprinting at the H19 – Igf2 locus maintains adult haematopoietic stem cell quiescence[J]. Nature, 2013, 500(7462): 345–349.

57. Jeong M, Sun D, Luo M, et al. Large conserved domains of low DNA methylation maintained by Dnmt3a[J]. Nature genetics, 2014, 46(1): 17–23.

58. Challen G A, Sun D, Jeong M, et al. Dnmt3a is essential for hematopoietic stem cell differentiation[J]. Nature genetics, 2012, 44(1): 23–31.

59. Zhang X, Jeong M, Huang X, et al. Large DNA methylation nadirs anchor chromatin loops maintaining hematopoietic stem cell identity[J]. Molecular cell, 2020, 78(3): 506–521. e6.

60. Kolmogorov M, Bickhart D M, Behsaz B, et al. metaFlye: scalable long-read metagenome assembly using repeat graphs[J]. Nature methods, 2020, 17(11): 1103–1110.

61. Durand N C, Shamim M S, Machol I, et al. Juicer provides a one-click system for analyzing loop-resolution Hi-C experiments[J]. Cell systems, 2016, 3(1): 95–98.

